# BNP facilitates NMB-mediated histaminergic itch via NPRC-NMBR crosstalk

**DOI:** 10.1101/2021.01.26.428310

**Authors:** Qing-Tao Meng, Xian-Yu Liu, Xue-Ting Liu, Devin M. Barry, Hua Jin, Yu Sun, Qianyi Yang, Li Wan, Jin-Hua Jin, Kai-Feng Shen, Admire Munanairi, Ray Kim, Jun Yin, Ailin Tao, Zhou-Feng Chen

## Abstract

B-type natriuretic peptide (BNP) binds to its two cognate receptors NPRA and NPRC, encoded by *Npr1* and *Npr3*, respectively, with equal potency and both are expressed in the spinal cord. Moreover, natriuretic peptides (NP) signal through the inhibitory cGMP pathway, raising the question of how BNP may transmit itch information. We report that *Npr3* is highly restricted to laminae I-II of the dorsal horn, and partially overlaps with neuromedin B receptor (NMBR) that encodes histaminergic itch. Functional studies indicate that NPRC is required for itch evoked by histamine but not chloroquine (CQ), a nonhistaminergic pruritogen. Importantly, BNP significantly facilitates scratching behaviors mediated by NMB, but not gastrin releasing peptide (GRP) that encodes nonhistaminergic itch. Consistently, BNP evoked Ca^2+^ response in NMBR/NPRC HEK 293 cells and BNP-saporin that ablated both *Npr1* and *Npr3* neurons impaired histamine-, but not CQ-evoked, itch. These results reveal a previously unknown mechanism by which BNP changes its inhibitory mode of action to the facilitation of itch through a novel NPRC-NMBR cross-talk. Our studies suggest that neuropeptides encode histaminergic and nonhistaminergic itch not only through distinct modes but also in synergy.

## Introduction

Neuropeptides are crucial for the transmission and modulation of itch sensation from the periphery to the spinal cord. In response to pruritogenic stimuli, a key event is the release of a cohort of neuropeptides from primary afferents to activate various G protein coupled receptors (GPCRs) in the spinal cord (1–4). Although acute itch is protective and mostly mediated by histaminergic mechanisms, chronic itch is a major debilitating condition associated with a wide spectrum of disorders, including skin, immune, nerve and systemic diseases due to resistance to antihistamines as well as a lack of effective therapeutics (5–7). Human and animal studies have shown that histaminergic and nonhistaminergic itch is transmitted through discrete molecular machinery and parallel primary afferent pathways (8–11). Despite the importance of nonhistaminergic itch in the etiology of chronic pruritus, how diverse neurotransmitters regulate histaminergic vs nonhistaminergic itch transmission is not well understood. We have shown that gastrin releasing peptide (GRP) and neuromedin B (NMB), two mammalian neuropeptides related to amphibian bombesin (12), encode nonhistaminergic itch and histaminergic itch, respectively (13–17). Moreover, murine GRP-GRPR signaling is important for the development of chronic itch, and itch transmission from primary afferents to the spinal cord is independent of presynaptic glutamatergic transmission under normal conditions (18–23).

B-type or brain natriuretic peptide (BNP) encoded by the gene *Nppb* has been implicated in itch at discrete regions, including the skin cells, sensory neurons and spinal cord (22, 24–26). Aside from BNP, the natriuretic peptide (NP) family also consists of atrial (ANP) and C-type natriuretic peptides (CNP) (27, 28). BNP binds to both NPRA and NPRC, encoded by *Npr1* and *Npr3*, with equality affinity, but not NPRB, while ANP also binds NPRA directly, resulting in the elevation of the second message cyclic GMP concentration (Figure 1A) (28, 29). Although NPRC is considered to function as a clearance receptor, it can also mediate guanylyl cyclase (GC)-coupled G_αi_ signaling under certain physiological conditions (30). BNP-NPRA signaling was originally proposed as an itch-specific pathway responsible for both histamine- and CQ-evoked itch via GRP-GRPR signaling (25, 31). However, functionally, GRP is required only for nonhistaminergic but not histamine itch (13, 16, 17), while recent studies have shown that BNP-NPRA signaling is required for histaminergic itch as well as chronic itch in mice which comprises histaminergic component (22, 24). Given that BNP can bind both NPRA and NPRC, two cognate receptors for BNP (Figure 1A), the relationship between NPRA/NPRC and NMBR/GRPR, as well as the role of NPRC in itch transmission remain undefined. Moreover, considering that the GC-cGMP signal transduction pathway of BNP is inhibitory and BNP-NPRA/NPRC signaling likely exerts inhibitory rather than excitatory function, analogous to G_αI_ protein coupled signaling, it is paradoxical that BNP would transmit rather than inhibiting itch information. In the present study, we have examined these open questions using a combination of RNA-scope ISH, genetic knockout (KO) mice, spinal siRNA knockdown, cell ablation, calcium imaging, pharmacological and optogenetic approaches.

**Figure 1.**
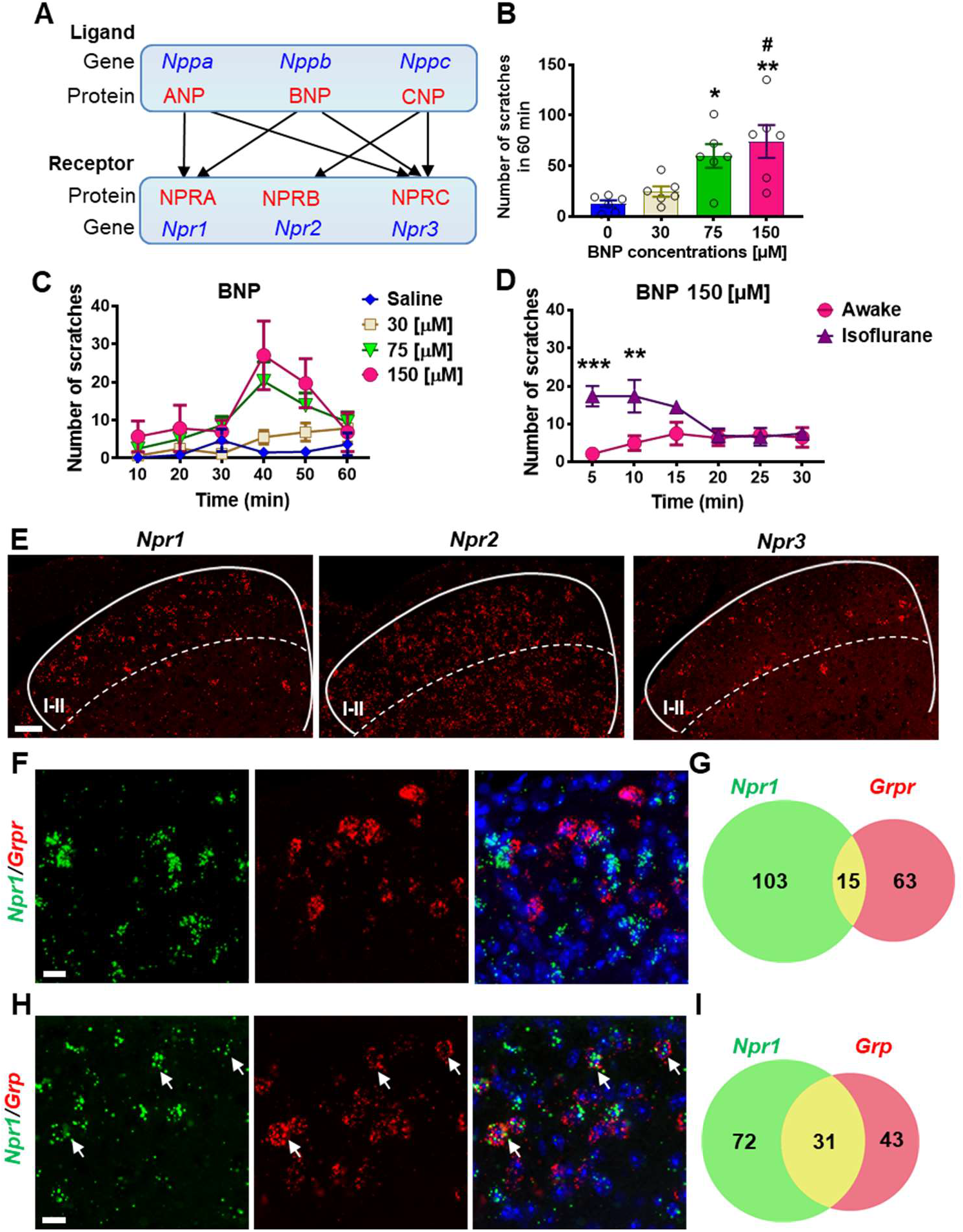
Expression of *Npr1*, 2 and 3 in the spinal cord. (**A**) A diagram shows crosstalk between NPs and NP receptors. BNP can bind NPRA and NPRC. (**B**) BNP dose-dependently evoked scratching behaviors 60 min after i.t. injection. n = 6. \**P* < 0.05, \*\**P* < 0.01, versus Saline, ^#^*P* < 0.05, versus 30 μM (equivalent to 1 μg), one-way ANOVA followed by Tukey’s test. (**C**) Time-course of scratching behaviors induced by different doses of BNP shows a delayed onset of scratching responses. (**D**) Scratching behaviors elicited by i.t. BNP (5 μg) were significantly enhanced by isoflurane. n = 6. \*\**P* <0.01, \*\*\**P* < 0.001, two-way ANOVA followed by Bonferroni's test. Values are presented as mean ± SEM. (**E**) Representative RNAscope ISH images show that *Npr1* and *Npr2* are widely expressed in the dorsal horn of the spinal cord, while *Npr3* is detected in the superficial dorsal horn. Scale bar, 100 μm. (**F**) Images of double ISH show that *Npr1* (green) does not overlap with *Grpr* (red) in the dorsal horn. Scale bar, 20 μm. (**G**) Venn diagram shows an overlap of *Npr1* and *Grpr* expression in the dorsal horn. (**H**) Images of double ISH show that *Npr1* (green) is co-expressed with *Grp* (red) in the dorsal horn. Arrows indicate double positive neurons. Scale bar, 20 μm. (**I**) Venn diagram shows an overlap of *Npr1* and *Grpr* expression in the dorsal horn.

## Results

### Expression of NP receptors in the spinal cord

As ANP also binds NPRA at high affinity (32, 33), we tested whether intrathecal injection (i.t.) of ANP could induce scratching behavior and found that ANP failed to induce scratching behaviors at the dose of 5 - 10 μg (data not shown). In contrast, BNP evoked dose-related scratching behavior at dose 1 – 5 μg (equivalent to 30-150 μM) with the peak scratching number of 74±16.2 (Figure 1B), consistent with previous reports (34, 35). However, these doses are much higher than endogenous concentrations of ligands that should be within the nanomolar range, implying a non-specific pharmacological effect. Time course analysis showed that scratching behavior was delayed by 20-30 min after BNP injections as described without isoflurane treatment (34, 35) (Figure 1C). Since some researchers performed i.t. injection on animals pre-treated with isoflurane for anesthesia and effects by isoflurane on neural circuits are extremely complex (36, 37), we compared the effect of BNP on awake animals and isoflurane-anesthetized animals and found that isoflurane pre-treatment significantly enhanced BNP-induced scratching behaviors in the first 10 min (Figure 1D), suggesting that induced scratching behavior is indirect rather than a direct effect of BNP with isoflurane treatment, consistent with the fact that NPRA/NPRC are inhibitory receptors. Recent studies show that *Npr1* is widespread in the dorsal horn (22, 38), in marked contrast to laminae I-II specific Grp expression (17, 39–41). Consistently, single nucleus RNA sequencing (snRNA-seq) from isolated spinal cord neurons found only partial overlap (20 ~ 30%) of *Npr1* and *Grp* (42). To visualize the distribution of NP receptors in the spinal cord, we performed *in situ* hybridization (ISH) using RNAscope probes, enabling highly sensitive and a specific detection of single transcript of target genes (43). Confocal images of RNAscope ISH showed that all three NP receptors are expressed in the dorsal horn of the spinal cord (Figure 1E). Consistent with a previous ISH study (22, 25, 38) and Allan Brain Atlas database (2), *Npr1*^+^ neurons distribute in a gradient manner with higher intensity throughout laminae I-IV and are rather sparse in the deep dorsal horn (Figure 1E). Remarkably, *Npr3*^+^ neurons are predominantly restricted to lamina I-II (Figure 1E). In contrast, *Npr2* expression is homogenous throughout the spinal cord dorsal horn, implying it has no modality-specific function (Figure 1E). There is a minimal overlapping expression between *Npr1* and *Grpr* (Figure 1F, G). However, approximately 30% (31/103) of *Npr1*^+^ neurons in laminae I-II of the dorsal horn express *Grp*, and this number is further reduced to 23% (31/132) (data not shown) when all *Npr1*^+^ neurons in the dorsal horn were counted (Figure 1H, I).

### NPRA and NPRC are important for acute itch

To examine the role of NP receptors in acute itch behavior, we first analyzed the phenotype of *Npr1* knockout (KO) mice (44). A previous study that found CNP-NPRB is essential for axonal bifurcation of DRG neurons in the developing spinal cord (45) prompted us to evaluate the innervation of primary afferents in the spinal cord of *Npr1* KO mice. We found that innervations of peptidergic CGRP^+^ and non-peptidergic IB4^+^ primary afferents in the superficial dorsal horn of *Npr1* KO mice are comparable with wild-type (WT) littermates (Supplemental Figure 1A and 1B). The innervations of GRP^+^, TRPV1^+^ and SP^+^ primary afferents are also comparable between WT and *Npr1* KO mice (Supplemental Figure 1C-1H), indicating that NPRA is dispensable for innervation of primary afferents.

GRP and NMB have been implicated in nonhistaminergic and histaminergic itch, respectively (13, 16, 46). To examine whether GRPR and NMBR function normally in the absence of NPRA, we compared the scratching behaviors between *Npr1* KO mice and their WT littermates after i.t. GRP or NMB and found no significant differences in their responses to either GRP or NMB between the groups (Figure 2A). However, *Npr1* KO mice showed significantly impaired scratching responses to intradermal (i.d.) injection of histamine and chloroquine (CQ), archetypal pruritogens for the histaminergic and nonhistaminergic itch, respectively (14), as compared with WT littermates (Figure 2B).

**Figure 2.**
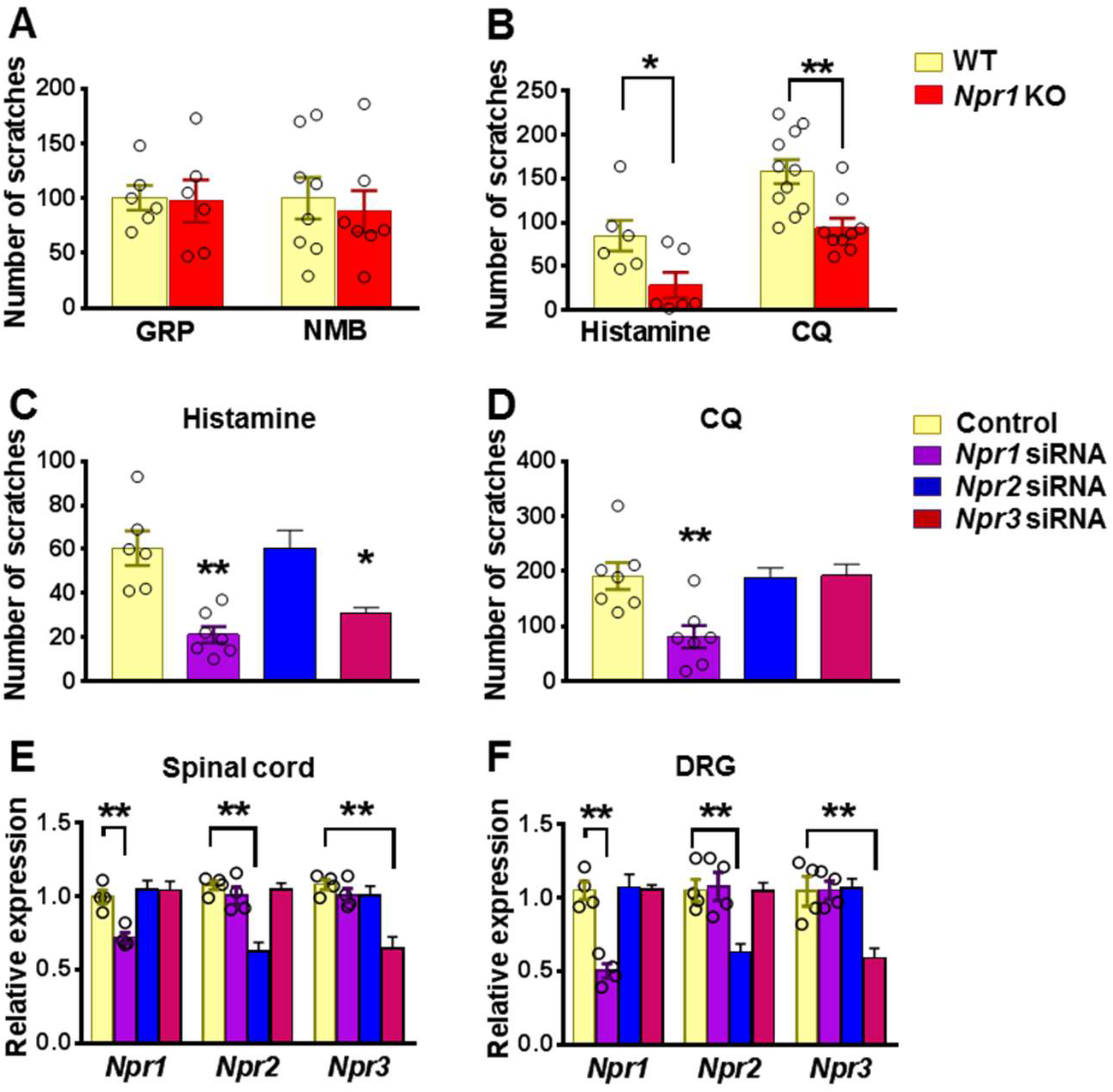
NPRA and NPRC are involved in acute itch. (**A**) *Npr1* KO mice and their WT littermates showed comparable scratching behaviors in response to GRP (5 μM, i.t.) and NMB (50 μM, i.t.). n = 6-8. (**B**) *Npr1* KO mice showed significantly reduced scratching behaviors elicited by histamine (200 μg, i.d.) and CQ (200 μg, i.d.). n = 9-11. \**P* < 0.05, \*\**P* < 0.01, unpaired t test. (**C** and **D**) Mice treated with i.t. *Npr1* siRNA and *Npr3* siRNA showed significantly reduced scratching responses to histamine (200 μg, i.d.) (**C**). CQ (200 μg, i.d.) itch was significantly reduced in mice that received *Npr1* siRNA treatment (**D**). n = 6-8. \**P* < 0.05, \*\**P* < 0.01, one-way ANOVA followed by Dunnett’s test. (**E** and **F**) Real-time PCR confirmed the reduced *Npr1-3* expression by siRNA knockdown targeting *Npr1*, *Npr2*, and *Npr3* in the spinal cord (**E**) and DRG (**F**), respectively. n = 4. \*\**P* < 0.01, one-way ANOVA followed by Dunnett’s test. Values are presented as mean ± SEM.

The highly restricted expression of NPRC in lamina II and its high binding affinity to BNP prompted us to examine the role of NPRC in itch. However, *Npr3* KO mice showed severe skeletal abnormalities, failing most KO mice to survive to the adult stage for behavioral analysis (47). To circumvent the problem, we performed spinal *Npr3* siRNA knockdown in mice. Moreover, to further determine whether the impaired scratching response of the global *Npr1* KO mice could have resulted from the *Npr1* deficiency in the spinal cord or DRGs where *Npr1* is also expressed (48), or the skin cells (26), we knocked down *Npr1-3* either individually or in combination in C57/BL6 mice using sequence specific siRNA. I.t. *Npr1* siRNA treatment significantly attenuated the scratching behavior evoked by histamine and CQ (Figure 2C and 2D), whereas *Npr3* siRNA treatment selectively attenuated histamine, but not CQ itch (Figure 2C and 2D). *Npr2* siRNA had no effect on CQ and histamine itch, making it unlikely to be involved in itch transmission (Figure 2C and 2D). The effect of knock-down of target mRNA in the spinal cord and DRGs was verified using real time RT-PCR (Figures 2E and 2F). These results revealed that *Npr1* and *Npr3* are differentially required for acute itch behavior at the spinal level. Further, we infer that there is little functional compensation among the three NP receptors.

### BNP facilitates NMB-mediated histamine itch

The slow onset of scratching behavior elicited by BNP, even at a high dose (150 μM), in the first 30 min contrasts sharply with the fast onset of GRP/NMB-induced scratching behavior, implying that direct activation of NPRA/NPRC alone is insufficient to initiate scratching response. This raised the question as to the specific role BNP may play in the first phase, which is physiologically relevant to histamine and CQ itch that usually occurs within this period. We suspected that BNP may play a modulatory function in acute itch behavior in a manner resembling the role of serotonin in itch modulation (49). To test this, we pre-treated mice with BNP at a lower dose (30 μM, i.t.) followed by i.d. histamine at the dose of 100 μg that is insufficient to induce robust scratching behaviors. At 30 μM, BNP failed to induce the scratching behaviors (Figure 1B). Strikingly, histamine-induced scratching responses were significantly enhanced with BNP pretreatment compared with those treated with saline (Figure 3A). BNP also produced a similar potentiating effect on CQ-induced scratching responses (Figure 3B). Since NMB is required for histamine itch via NMBR exclusively (13, 16), we tested the possibility that BNP may facilitate histamine itch by modulating NMBR function. At 5 μM, i.t. NMB alone could not induce significant scratching behavior (Figure 3C). However, co-injection of BNP (30 μM) and NMB (5 μM) markedly increased NMB-induced scratching behavior compared with that of mice receiving only NMB (Figure 3C). Importantly, BNP failed to potentiate scratching behaviors induced by GRP (1 μM, i.t.), which is not required for histamine itch (Figure 3D).

**Figure 3.**
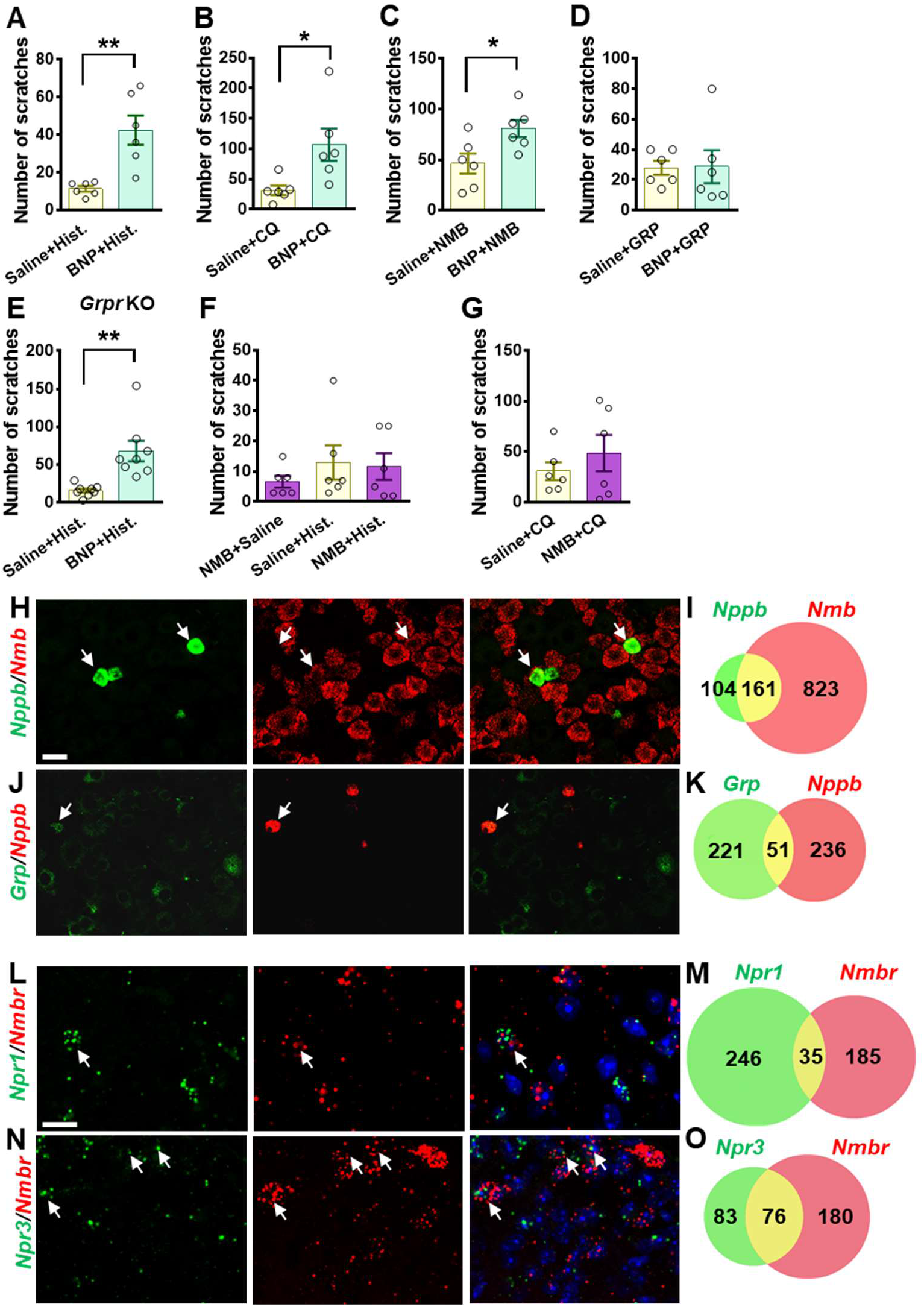
BNP facilitates histamine itch. (**A**) Pre-injection of BNP (30 μM, i.t.) for 1 min significantly enhanced scratching behavior evoked by i.d. injection of histamine (Hist.) (100 μg). n = 6. (**B**) Scratching behavior evoked by i.d. injection of CQ (50 μg, i.d.) was significantly enhanced by pre-injection of BNP for 1 min. n = 6. (C and D) Co-injection of BNP (30 μM, i.t.) facilitated scratching behaviors evoked by NMB (5 μM, i.t.) (**C**) but not GRP (1 μM) (D). n = 6. (**E**) Pre-injection of BNP (30 μM, i.t.) for 1 mine significantly enhanced scratching behavior evoked by i.d. injection of histamine (100 μg) in *Grpr* KO mice. n = 8. (**F and G**) Pre-injection of NMB (5 μM, i.t.) had no effect on scratching behaviors induced by histamine (**F**) or CQ (**G**). Note that NMB only evoked a minimal number of scratching bouts towards the nape area where histamine and CQ were intradermally injected. n = 6. (**H-K**) Double RNAScope ISH images (**H** and **J**) and Venn diagrams (**I** and **K**) show 60% of *Nppb* neurons co-expression *Nmb* in DRG (**H** and **I**), while *Grp* neurons show minimal co-expression with *Nppb* (**J** and **K**). Scale bar, 20 μm. (**L-O**) Double RNAScope ISH images (**L** and **N**) and Venn diagrams (**M** and **O**) show minimal co-expression of *Npr1* and *Nmbr* in the superficial dorsal horn (**L** and **M**), while 48% of *Npr3*^+^ neurons also express *Nmbr* (**N** and **O**). Scale bar, 50 μm. Values are presented as mean ± SEM, \**P* < 0.05, \*\**P* < 0.01, unpaired t test in (A-E), one-way ANOVA in (F and G). Scale bar, 50 μm.

We then assessed whether BNP may function upstream or independent of GRPR to modulate itch by comparing the facilitatory effect of BNP on histamine itch between *Grpr* KO and WT mice. If BNP acts upstream of or depends on GRPR, BNP may fail to potentiate histamine itch in *Grpr* KO mice. Indeed, we found that BNP similarly potentiated histamine itch in *Grpr* KO mice (Figure 3E), consistent with the findings that GRP-GRPR signaling is not required for histamine itch (46). Together, these results show that the role of BNP signaling in the spinal cord is dependent on NMB-NMBR signaling and independent of GRP-GRPR signaling.

Next, we evaluated whether NMB has a modulatory function that resembles BNP in histamine itch. We found that mice pretreated with NMB (5 μM, i.t.) failed to exhibit enhanced scratching behaviors evoked by either histamine or CQ (Figures 3F and 3G), suggesting that it is unlikely that NMB would function as a modulator. We previously showed that NMB exerts its role exclusively through NMBR in the spinal cord, as NMB is a functional antagonist for GRPR in spite of its cross-binding activity with GRPR (13). This prompted us to test whether BNP may facilitate NMB/histamine itch signaling through crosstalk between NMBR, which is required for histamine itch, and NPRA or NPRC, two receptors that bind to BNP. Double RNAscope ISH showed that ~60% of *Nppb* neurons co-expressed *Nmb* in the DRG (Figures 3H and 3I), whereas *Grp* neurons in DRG showed little co-expression (~19%) with *Nppb* (Figures 3J and 3K). Moreover, double ISH showed that *Npr1* and *Nmbr* minimally overlap in the dorsal horn (Figures 3L and 3M), excluding the likelihood of NPRA-NMBR cross-signaling. By contrast, approximately 47.8% of *Npr3*^+^ neurons express *Nmbr* (Figures 3N and 3O), raising the possibility that NPRC is involved in crosstalk with NMBR.

### BNP facilitates NMB-evoked calcium response via NPRC-NMBR cross-signaling

To test whether BNP can potentiate NMBR function, we took advantage of the fact that NMB exclusively activates NMBR neurons in the spinal cord (13) and examined the response of NMBR neurons to NMB using Ca^2+^ imaging of dorsal horn neurons (50). NMBR functions via the canonic G_q_ coupled PLC-PKC-Ca^2+^ signaling similar to GRPR (49, 51). Using a protocol for investigating facilitating effect (49), we found that NMB at 20 nM was able to induce Ca^2+^ transient, but not at 10 nM, in perspective NMBR neurons (identified with the first application) (Figure 4A). BNP alone could not induce Ca^2+^ transient, regardless of the dose, in these neurons. Strikingly, when BNP (200 nM) was co-applied with the subthreshold concentration of NMB (10 nM), it dramatically potentiated Ca^2+^ transients in response to the second NMB application (Figure 4A). Importantly, NMB (up to 20 nM) failed to induce Ca^2+^ transients in the dorsal horn neurons isolated from *Nmbr* KO mice, further demonstrating the specificity of the response of NMBR neurons to NMB (Figure 4B).

**Figure 4.**
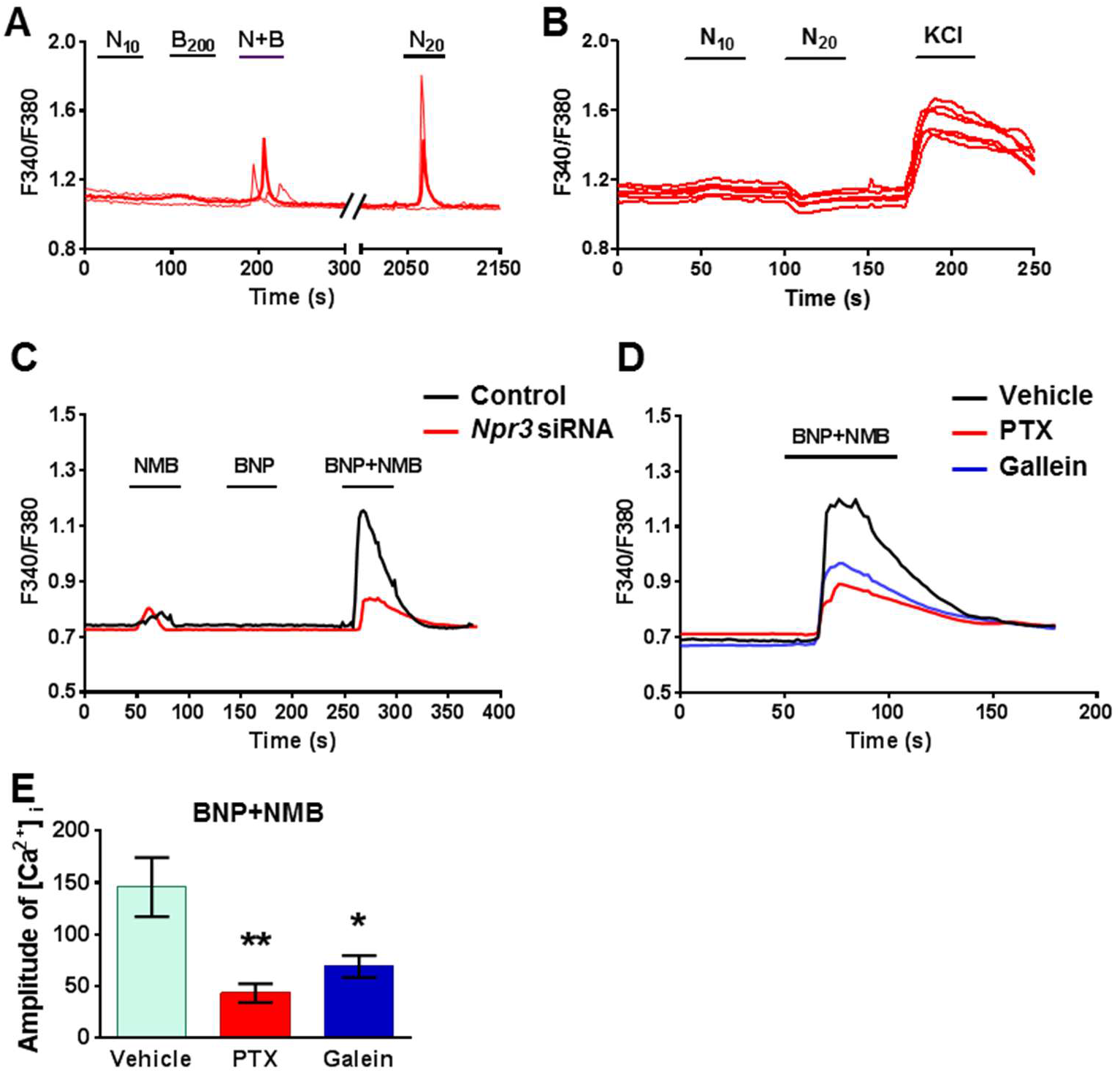
BNP facilitates calcium response induced by NMB in spinal cord neurons and HEK293 cells. (**A**) Co-application of BNP (200 nM)(B_200_) facilitates the calcium response of dorsal horn neurons to NMB (10 nM)(N_10_). (**B**) NMB up to 20 nM (N_20_) did not evoke calcium response on dorsal horn neurons of *Nmbr* KO mice. (**C**) Co-application of BNP (1 μM) with subthreshold of NMB (1 pM) evoked robust calcium response in HEK 293 cells expressing NMBR. (**D**) Calcium responses induced by BNP+NMB were attenuated by pre-incubation of PTX (200 ng/ml) or gallein (100 μM). (**E**) Quantification of intracellular calcium concentration ([Ca^2+^]_i_) shows that PTX and gallein significantly blocked the calcium release induced by BNP+NMB. Values are presented as mean ± SEM, n = 6. \**P* < 0.05, \*\**P* < 0.01, one-way ANOVA followed by Tukey’s test.

To probe the possibility of NPRC-NMBR crosstalk, we took advantage of the finding that HEK 293 cells express endogenous *Npr1* and *Npr3* as shown by qRT-PCR (Supplemental Figure 2). In HEK 293 cells stably expressing NMBR, we co-applied BNP (1 μM) and NMB (1 pM), which induces minimal Ca^2+^ response when applied individually, and observed robust Ca^2+^ transients (Figure 4C). Importantly, the effect of BNP was greatly attenuated by *Npr3* siRNA treatment (Figure 4C), indicating that BNP facilitates NMB/NMBR signaling through NPRC. NPRC has been linked to the inhibition of adenylate cyclase (AC)/cAMP signaling and can activate the pertussis toxin (PTX)-sensitive G_αi/βγ_ signaling pathway (52–54). To examine whether the G_αi/βγ_ pathway is involved in the facilitatory effect of BNP/NPRC, we pre-incubated HEK 293 cells with pertussis toxin (PTX) to inactivate G_αi_ protein (55). Subsequent incubation of BNP and NMB following PTX treatment induced much smaller Ca^2+^ spikes compared to control cells (Figure 4D). The amplitude of intracellular Ca^2+^ concentrations ([Ca^2+^]_i_) was significantly reduced by PTX treatment (Figure 4E). Pre-incubation of gallein, a small molecule G_βγ_ inhibitor (56), also blocked the facilitation effect of BNP on NMB-induced calcium spikes (Figures 4D and 4E). These results suggest that BNP-NPRC signaling positively regulates NMB/NMBR signaling via G_αi/βγ_ signaling.

### NPRA and NPRC neurons are required for histamine itch

BNP-saporin (BNP-sap) has been used to ablate NPRA neurons in the spinal cord (25). Nevertheless, the expression of NPRC in the dorsal horn raised the question of whether BNP-sap may additionally ablate neurons expressing NPRC (Figure 1A). A premise for ablation of neurons with peptide-conjugated saporin approach is the internalization of the receptor upon binding to the saporin, resulting in cell death (57). To test whether BNP can also internalize NPRB and NPRC, HEK 293 cells were transfected with *Npr1, 2, 3* cDNA tagged with mCherry (mCh) separately. Consistent with previous studies (58, 59), BNP internalized NPRA or NPRC but not NPRB in HEK 293 cells (Supplemental Figure 3A), indicating that BNP-sap could ablate both NPRA and NPRC cells. Because BNP-sap at 5 μg as described previously (25) resulted in the lethality of C57 mice in our tests, we reduced the dose to 2.5 μg so that enough animals could survive for behavioral and molecular analysis. RNAscope ISH showed the number of *Npr1*^+^ neurons were reduced to ~50% (82.9 ± 3.3 in Control vs. 39.8 ± 2.2 in BNP-sap) (Figures 5A and 5B), whereas *Npr2*^+^ neurons were not affected (288.0 ± 18.2 in Control vs. 300.5 ± 7.8 in BNP-sap) (Figures 5C and 5D). Moreover, BNP-sap ablated ~67% of *Npr3*^+^ neurons (41.8 ± 1.4 in Control vs. 13.8 ± 1.8 in BNP-sap) as well as ~37% of Grp^+^ neurons (44.6 ± 1.9 in Control vs. 22.6 ± 2.0 in BNP-sap) (Figures 5G and 5H). As expected, the number of *Nmbr*^+^ neurons were also significantly reduced after BNP-sap injection, likely due to *Npr3* expression in these neurons (Figures 5I and 5J). Real time RT-PCR analysis confirmed reduced *Npr1*, *Npr3*, and *Nmbr* mRNA levels in the dorsal horn by BNP-sap treatment, whereas the expression of *Npr2* and Grpr was unaffected (Supplemental Figure 3B). Interestingly, behavioral studies showed that histamine itch was significantly reduced in BNP-sap mice (Figure 5K), whereas CQ itch was not affected (Figure 5L). These results suggest that NPRA and NPRC neurons in the spinal cord play an important role in histamine itch, which can be attributed to partial ablation of NMBR neurons.

**Figure 5.**
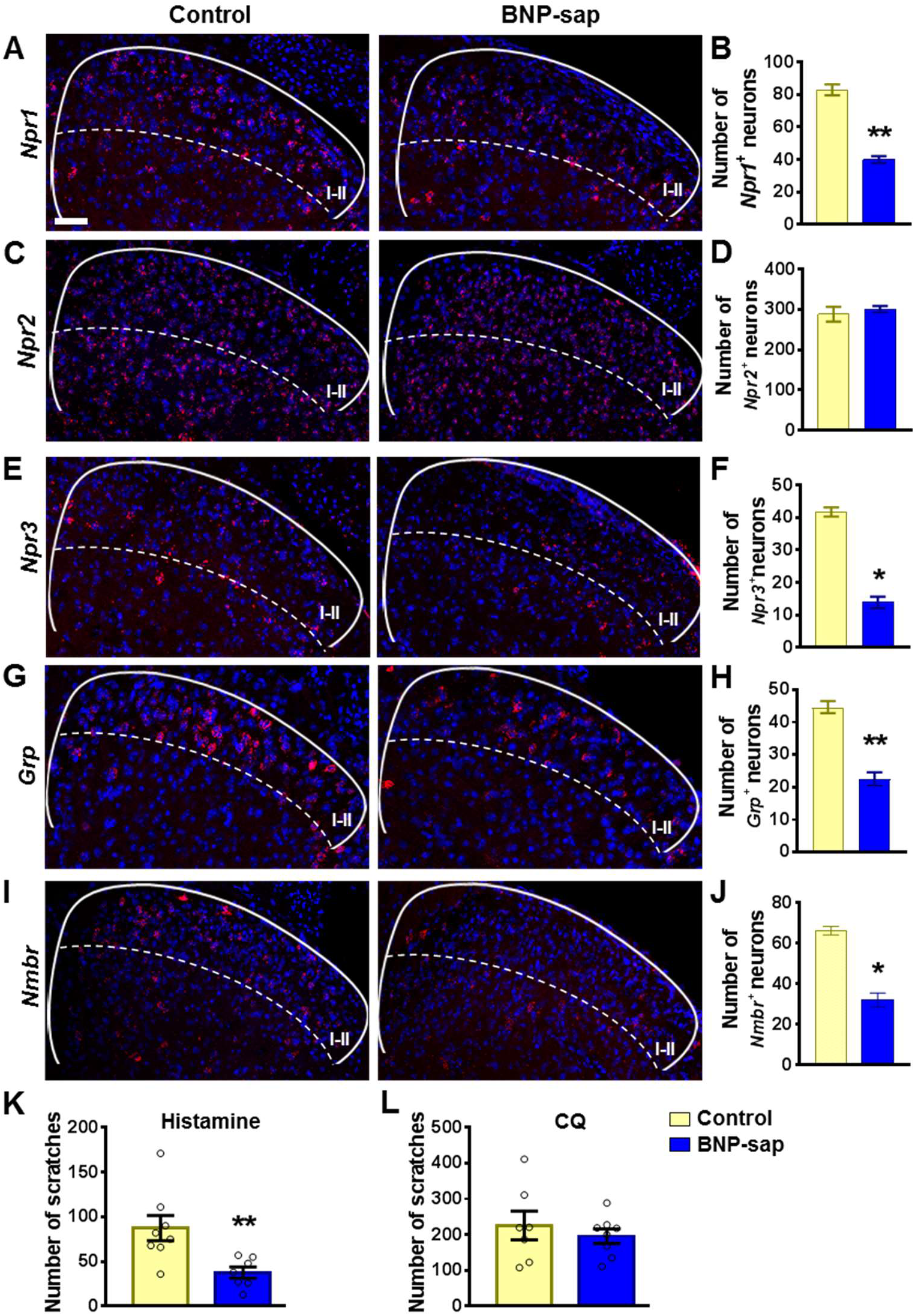
BNP-sap ablates spinal cord neurons expressing *Npr1* and *Npr3*. (**A-F**) RNAscope ISH images (**A**, **C**, and **E**) and quantification (**B**, **D**, and **F**) show that *Npr1*^+^ (**A** and **B**) and *Npr3*^+^ (**E** and **F**) neurons in the dorsal horn of the spinal cord are ablated by BNP-sap, while *Npr2*^+^ (**C** and **D**) neurons are not affected. n = 4. (**G-J**) RNA scope ISH images (**G** and **I**) and quantification (**H** and **J**) show that the number of *Grp*^+^ neurons (**G** and **H**) and *Nmbr*^+^ neurons (**I** and **J**) are reduced in the superficial dorsal horn of BNP-sap mice. n = 4. (**K** and **L**) Scratching behaviors induced by histamine (**K**), but not CQ (**L**) were significantly reduced in BNP-sap treated mice. n = 7-8. Values are presented as mean ± SEM. \**P* < 0.05, \*\**P* < 0.01, unpaired t test. Scale bar, 50 μm.

### NPRA is dispensable for the development of dry skin itch and a mouse model of neuropathic itch

Finally, we assessed whether NPRA is important for chronic itch transmission by comparing spontaneous scratching behaviors between *Npr1* KO and WT littermate mice using a mouse model of dry skin (60). From day 8 *Npr1* KO mice appeared to show a tendency toward reduced scratching behavior relative to their WT littermates, the differences, however, were not statistically significant (Figure 6A). The normal dry skin itch in *Npr1* KO mice prompted us to examine BNP expression in DRGs of BRAF^Nav1.8^ mice that developed spontaneous scratching behavior, likely reflecting neuropathic itch resulting from enhanced expression of itch-sensing peptides/receptors in sensory neurons (21). To our surprise, real time RT-PCR results showed that the levels of *Nppb* mRNA in BRAF^Nav1.8^ mice, which developed spontaneous scratching behavior due to enhanced GRP expression in DRGs (21), were significantly reduced relative to the control mice (Figure 6B). Interestingly, BRAF^Nav1.8^ mice also showed decreased expression of somatostatin (*Sst*), a peptide co-localized with *Nppb* in DRG neurons (31, 61). Double ISH using RNAscope showed that about 82.5% of *Nppb*^+^ neurons and 74.0% of *Sst*^+^ neurons overlapped in DRGs (Figures 6C and 6D). Consistent with PCR results, *Nppb* signals were very weak, whereas Sst signals were barely detectable in DRGs of BRAF^Nav1.8^ mice (Figures 6C and 6E). In addition, conventional ISH showed that expression of *Nppb* was significantly reduced in DRGs of BRAF^Nav1.8^ mice (Supplemental Figure 4). Furthermore, we employed RNA-seq to compare gene expression profiles between BRAF^Nav1.8^ mice and WT mice. In line with previous findings (21), transcript levels of *Grp* and *Mrgpra3* were dramatically increased, whereas *Nppb* and *Sst* transcripts were markedly reduced in DRGs of BRAF^Nav1.8^ mice (Table 1). Taken together, these findings suggest that BNP-NPRA signaling is not required for the development of dry skin itch and neuropathic itch of BRAF^Nav1.8^ mice.

**Table 1.**
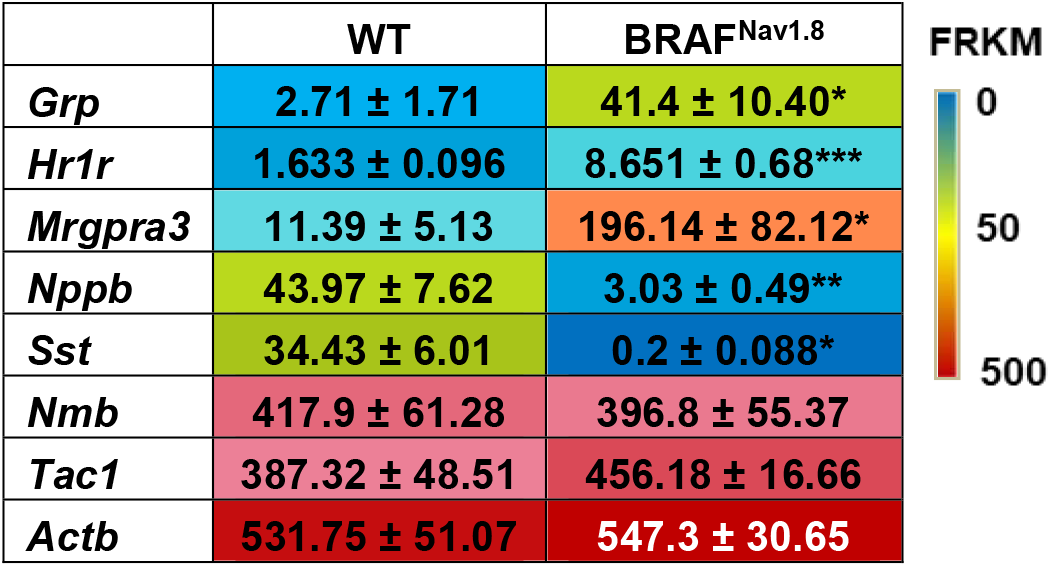
Gene expression in DRGs of WT and BRAF_Nav1.8_ mice.

**Figure 6.**
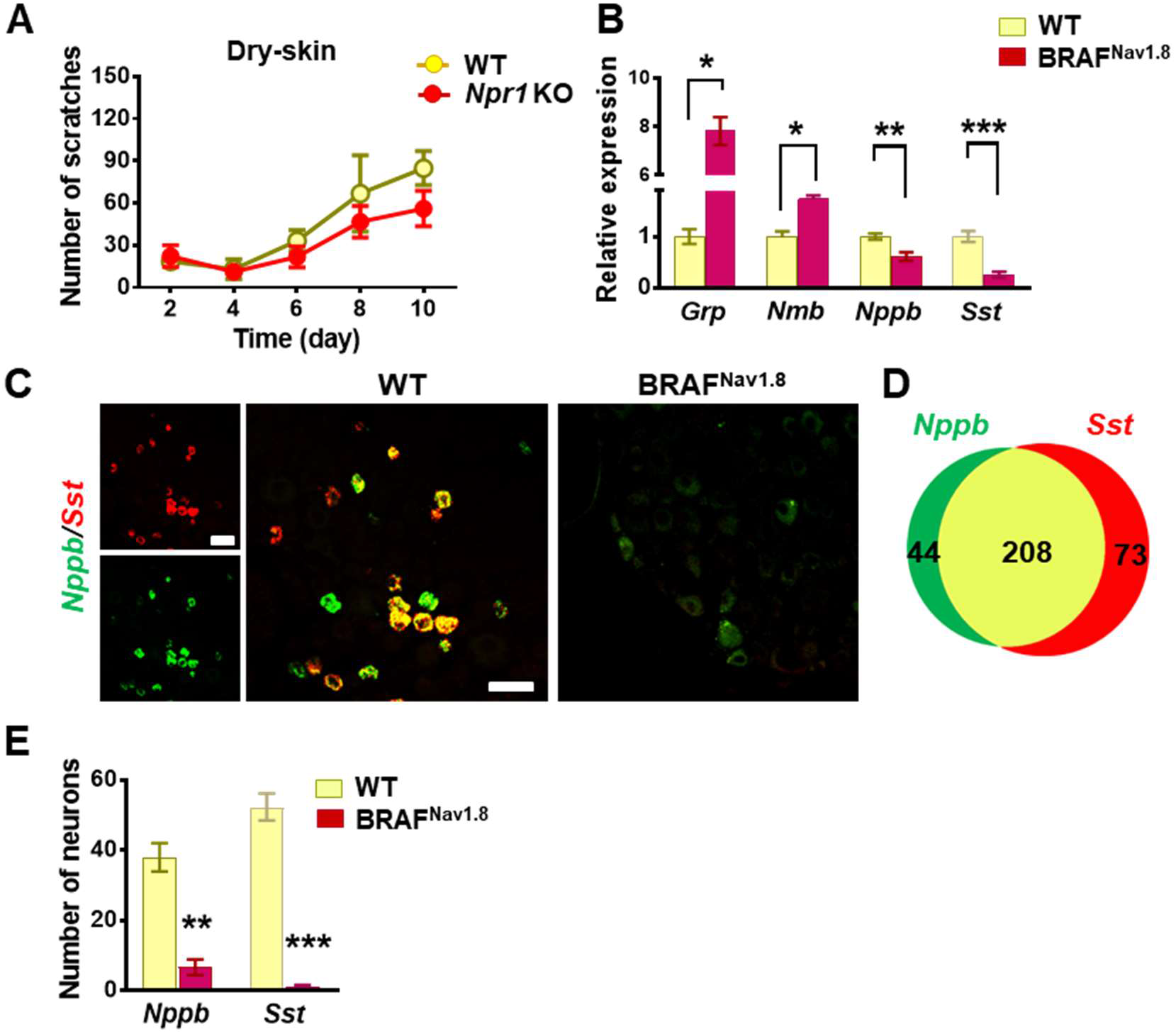
BNP-NPRA signaling is dispensable for chronic itch. (**A**) *Npr1* KO mice and WT littermates showed comparable spontaneous scratching behaviors in the dry skin model. n = 6, p = 0.1283, *F*_1,50_ = 2.392, repeated measures Two-way ANOVA. (**B**) Real-time RT PCR shows that the levels of *Nppb* mRNA and *Sst* mRNA in DRGs of BRAF^Nav1.8^ mice were significantly reduced compared with WT mice. n = 4. (**C**) RNA scope ISH images show that *Nppb* and *Sst* are largely co-expressed in WT DRG neurons. *Nppb* and Sst signals are dramatically reduced in DRGs of BRAF^Nav1.8^ mice. (**D**) Venn diagram shows an overlap of *Nppb* and *Sst* expression in WT DRGs. (**E**) Quantified data of RNA scope shows that the number of *Nppb*^+^ neurons and *Sst*^+^ neurons is significantly reduced in the DRGs of BRAF^Nav1.8^ mice. n = 4. Values are presented as mean ± SEM. \**P* < 0.05, \*\**P* < 0.01, \*\*\**P* < 0.001, unpaired t test. Scale bars, 50 μm.

Sst type 2 receptor (SST2R) is expressed in GABAergic neurons in the spinal cord and has been considered to be a sole receptor for SST (62, 63). The reduced Sst expression in chronic itch conditions may suggest a dampening of the dorsal horn GABAergic neuronal activity to enable enhanced itch transmission. To test whether SST2R neurons may inhibit both itch and pain transmission, we pharmacologically activated these neurons by i.t. SST followed by evaluating the nature of evoked scratching behavior since the injection onto the nape may induce itch, pain-related or undefined scratching behavior (64). We found that i.t. SST-evoked scratching was markedly reduced but not abolished by intraperitoneal injection (i.p.) of morphine (Figure 7A), a method used to evaluate whether i.t. induced scratching/biting behavior reflects pain (65). In addition, scratching behaviors evoked by i.t. injection of SST and octreotide (OCT), a selective SST2R agonist, were significantly attenuated on mice with bombesin-saporin (BB-sap) treatment, which can completely block nonhistaminergic itch transmission (46)(Figure 7B and 7C). These results suggest that pharmacological activation of SST2R neurons could result in disinhibition of both itch and pain transmission.

**Figure 7.**
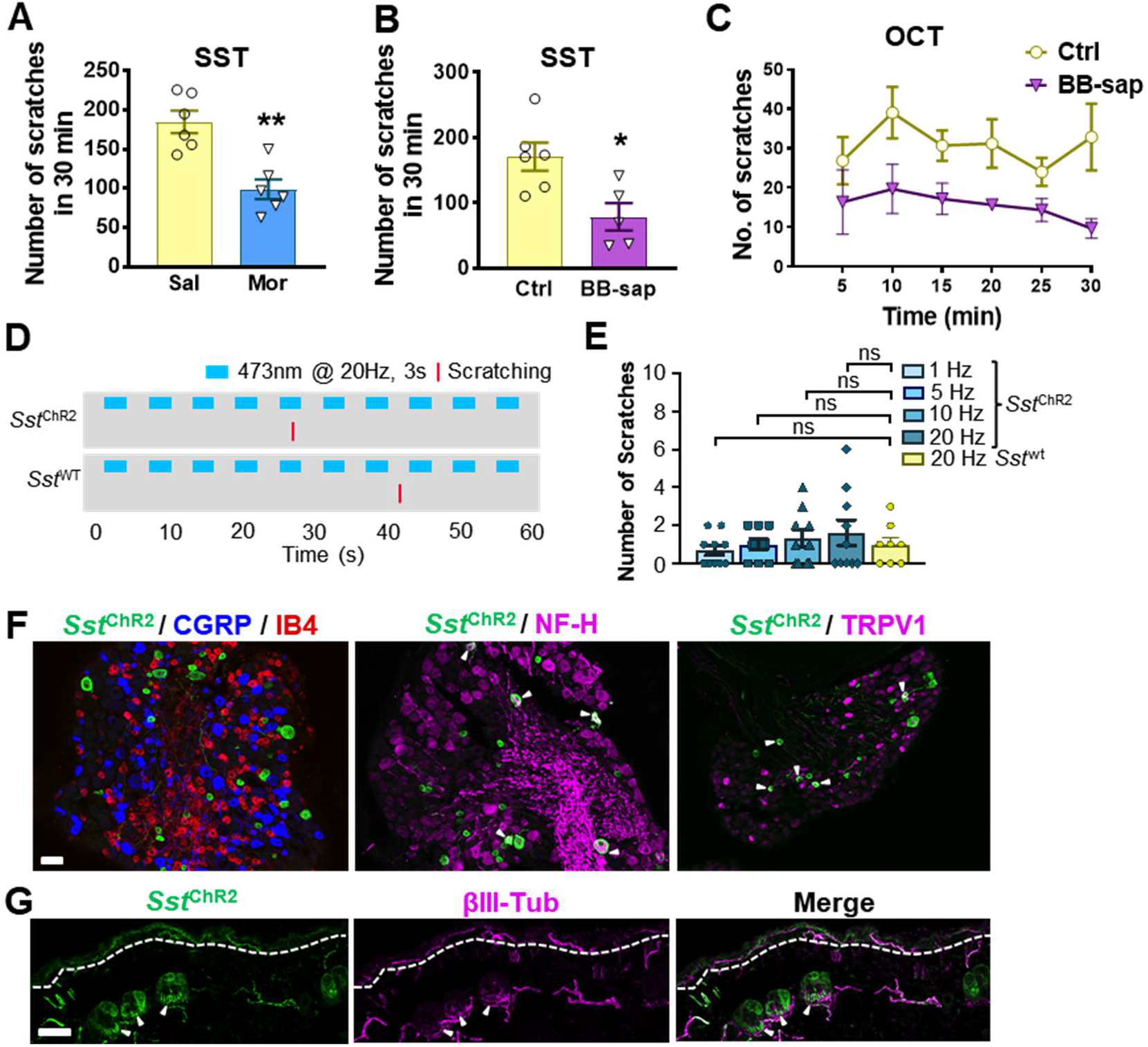
SST evoked both pain and itch responses in mice. (**A**) Pre-injection of morphine (10 mg/kg, i.p.) for 30 min attenuated scratching behaviors induced by i.t. injection of SST (5 nmol). n = 6 mice per group. Sal, saline; Mor, morphine. (**B** and **C**) SST (5 nmol, i.t.)(B) and OCT (C) -evoked scratching behaviors were significantly reduced in bombesin-saporin-treated mice comparing with control mice that were treated with blank saporin. n = 5-6 mice per group. Ctrl, control; BB-sap, bombesin-saporin. (**D**) Raster plot of scratching behavior induced by light stimulation of the skin in *Sst*^ChR2^ and *Sst*^WT^ mice. (**E**) The number of scratches in 5 min induced by 3s – 1, 5, 10 or 20 Hz light stimulation of nape skin in *Sst*^ChR2^ and *Sst*^WT^ mice. n = 8 - 10 mice. ns – not significant, one-way ANOVA with Tukey post hoc. (**F**) IHC images of *Sst*^ChR2^/CGRP/IB4 (left), *Sst*^ChR2^/NF-H (middle), and *Sst*^ChR2^/TRPV1 (right) in DRG of *Sst*^ChR2^ mice. Arrowheads indicate co-expression. (**G**) IHC image of *Sst*^ChR2^/βIII-Tubulin in hairy nape skin. The dashed line marks epidermal/dermal boundary. Arrowheads indicate ChR2 expression in lanceolate endings of hair follicles. Values are presented as mean ± SEM. \**P* < 0.05, \*\**P* < 0.01, unpaired t test. Scale bars, 100 μm.

**Figure 8.**
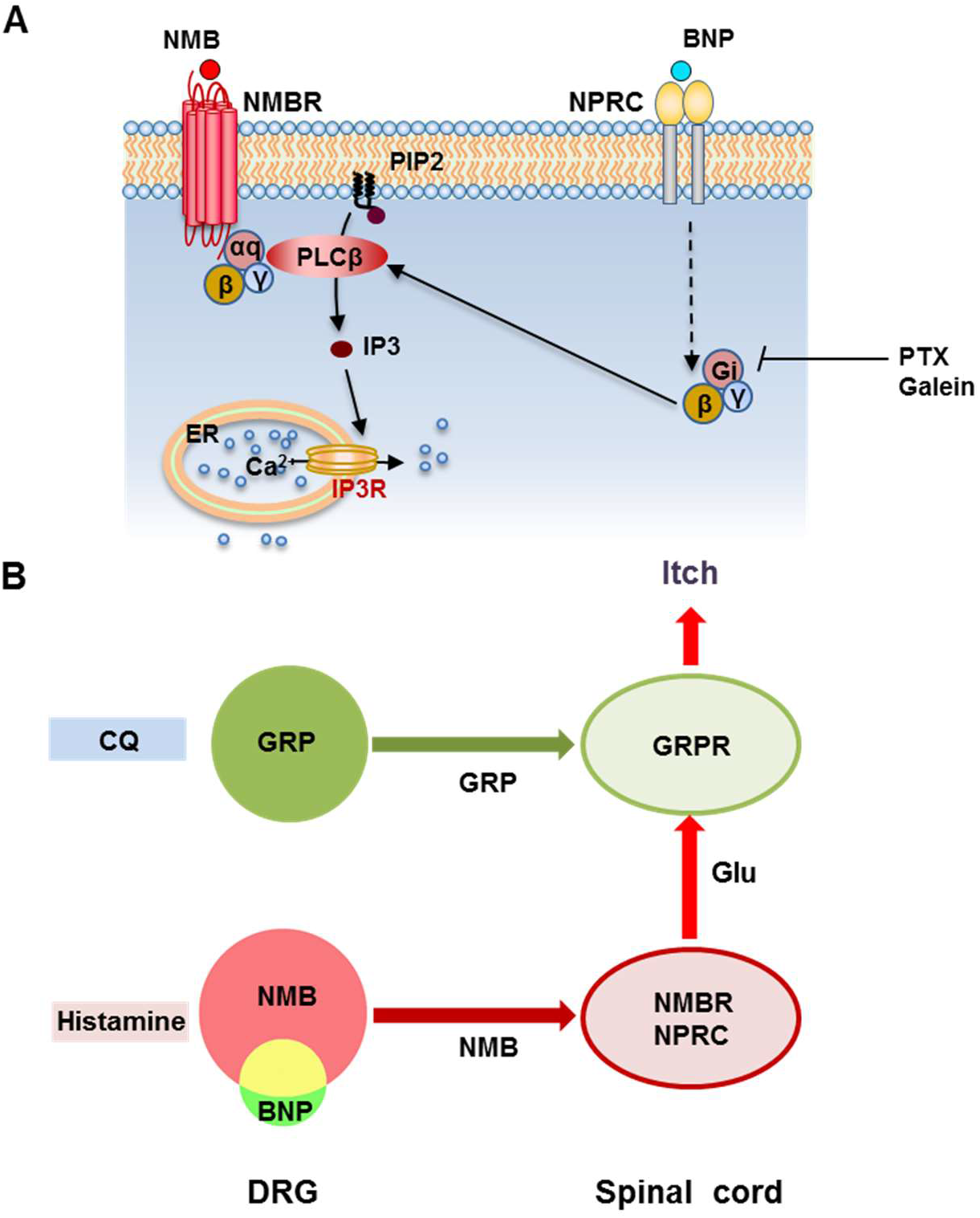
(**A**) A schematic showing a model for NMBR-NPRC cross-signaling facilitated by BNP via the NMB-NMBR pathway. In response to histamine, NMB and BNP are released from primary afferents to activate NMBR and NPRC concurrently. Activation of NMBR by NMB at a low concentration may prime PLC_b_ signaling, whereas activation of NPRC by BNP stimulates G_ai_ signaling, which in turn stimulates PLC_b_ to activate downstream Ca^2+^ signaling. (**B**) A hypothetic model depicting the respective roles of neuropeptides and glutamate in itch transmission. CQ itch is mediated in part by GRP-GRPR signaling independent of glutamatergic transmission. In contrast, histamine itch is mediated by NMB-NMBR signaling from primary afferents to NMBR neurons and by glutamatergic transmission from NMBR neurons to GRPR neurons. BNP facilitates NMB-NMBR signaling via NPRC independent of GRP-GRPR signaling but dependent on GRPR neurons. Glu: glutamate.

Optical activation of the skin innervated by *Grp* primary afferents evoked frequency-dependent itch-related scratching behavior (17). Given the largely overlapping of *Nppb* and *Sst* expression in DRGs, we used *Sst*-Cre mice as a surrogate for BNP-expressing fibers and examined whether optical stimulation of *Sst* expressing skin fibers is sufficient to evoke scratching behavior in mice. *Sst*^ChR2^ or *Sst*^WT^ mice were stimulated with 473 nm blue light with a fiber optic held just above the nape skin (15 mW power from fiber tip) at 1, 5, 10 or 20 Hz with a 3 s On-Off cycle for 5 min total (Figure 7D). Stimulation at all frequencies failed to evoke significant scratching behaviors in *Sst*^ChR2^ mice compared to *Sst*^WT mice^ (Figure 7E). At last, we analyzed the expression of *Sst*^ChR2^ in DRG. Consistent with previous studies (61), we found that *Sst*^ChR2^ sensory neurons do not co-express the peptidergic marker calcitonin gene-related peptide (CGRP), nor do they show the non-peptidergic Isolectin B4 (IB4)-binding (Figure 7F). However, some *Sst*^ChR2^ sensory neurons do co-express the myelinated marker neurofilament heavy (NF-H) as well as some co-expression with transient receptor potential cation channel subfamily V member 1 (TRPV1) (Figure 7F, arrowheads). Examination of the hairy nape skin revealed that expression of *Sst*^ChR2^ in the epidermis as well as expression in some hair follicles of the dermis within apparent lanceolate endings (Figure 7G, arrowheads).

The findings that *Sst* fibers innervate hair follicles (61), whereas GRP or *Mrgpra3* fibers only innervate the skin as free-nerve or bush/cluster endings mostly in the epidermis (17, 66, 67) supports organizational segregation of two distinct subpopulations of primary afferents in sensory neurons.

## Discussion

### The Role of Spinal NPRA and NPRA Neurons in Itch Transmission

The present study confirmed only a small subset of *Grp* neurons express *Npr1* (17) (38, 42), in agreement with conventional ISH results and ISH data from Allan Brain Atlas (2, 25). Although the siRNA knockdown of *Npr1* mRNA in both spinal cord and DRGs makes it difficult to ascribe deficits in itch to either region, the finding that BNP-sap treatment failed to impair CQ itch suggests that NPRA in DRGs rather than spinal cord is required for CQ itch. This interpretation is consistent with the observation that BNP did not facilitate GRP-induced scratching behavior, as well as the widespread expression of *Npr1* in the dorsal horn, suggesting that spinal NPRA is unlikely to have a modality-specific role in itch. Importantly, the finding that most of *Nppb* neurons express NMB, but not GRP, in DRGs implies that BNP/NMB and GRP released from distinct types of primary afferents target the dorsal horn neurons expressing NPRA/NPRC/NMBR and GRPR, respectively, matching organization segregation of SST/BNP and GRP afferents in the skin. Previous studies have shown that BNP/NPRA expression is upregulated in DRG neurons under inflammatory pain conditions and i.t. BNP reduced inflammatory pain (48). The failure of i.d. BNP and opto-stimulation of cutaneous STT/BNP fibers to evoke scratching behavior is in favor of the notion that NPRA in DRGs may facilitate CQ itch. To delineate a precise role of NPRA and its relation to NPRC in sensory neurons, generation of conditional knockout of *Npr1* in DRGs without affecting its expression in the spinal cord and skin is necessary.

Consistent with that BNP binds to NPRA/NPRC but not NPRB (28)(Figure 1A), we found that BNP internalizes NPRA/NPRC, but not NPRB. Accordingly, BNP-sap ablated NPRA and NPRC neurons in the spinal cord only without impacting NPRB neurons. Unlike BB-sap that only ablates GRPR neurons (46), one methodological limitation is that a high dose of BNP-sap may have ablated neurons expressing NP receptors in the brain, resulting in early death of mice. Therefore, normal CQ itch may be due to partial ablation of spinal NPRA/NPRC neurons. However, because mice treated with BNP-sap showed prominent deficits in histamine itch, a compensatory effect for CQ itch appears less likely. In addition, the observation that BNP did not modulate GRP-GRPR signaling also argues that NPRA and NPRC in the spinal cord are dispensable for CQ itch.

### The Modulatory Role of BNP-NPRC Signaling in Histamine Itch

The time-course analysis suggests that scratching behavior evoked by BNP is unlikely to have direct physiological relevance to itch transmission because BNP-evoked scratching showed a delayed onset, occurring only at relatively higher doses (2.5 - 5 μg), which likely falls out of the range of its endogenous concentration required for mediating acute itch transmission. By contrast, the potency of GRP or NMB in inducing scratching behavior is much higher than that of BNP (13, 14, 68). Although not feasible to measure in the mouse spinal cord, conceivably, the amount of GRP/NMB released endogenously in response to pruritogenic stimuli is likely to be a picomolar range, given their potent scratching-induced effect. In general, the lower dose of NMB used, the more likely it resembles physiological condition. That BNP potently enhances NMB function but not vice versa suggests an important mechanistic difference in their mode of action in the coding of itch information. In comparison, while glutamatergic signaling is required for itch transmission between NMBR and GRPR neurons (16) and between Tac2 neurons and GRPR neurons (23), it is dispensable from the periphery to the spinal cord (18, 19). This underscores the importance of neuropeptide signaling in the coding of itch information in sensory neurons (Figure 7B).

The most striking finding is that BNP-NPRC signaling plays an important role in the facilitation of histamine itch via NPRC-NMBR crosstalk. Earlier studies suggested that NPRC activation attenuates guanylyl cyclase/cAMP signaling, which in turn activates PLC signaling to stimulate IP3 production (30, 69), thereby hinting at a crosstalk between NP receptors and GPCRs. In cardiac cells where BNP and ANP, two important heart hormones, play a major role in the regulation of blood pressure, cGMP signaling has a facilitatory function by inhibiting the cAMP hydrolysis downstream of G_i_ protein coupled signaling (70). cGMP-mediated inhibition of cAMP signaling, possibly in a spatially defined cellular complex, appears to be a common mechanism by which NPs regulate a variety of physiological functions (70). Our finding that PTX/Galein inhibits the facilitatory effect of BNP on NMB-induced Ca^2+^ response in NPRC/NMBR cells suggests an intracellular coupling of the G_αi/βγ_ signaling pathway to PLC signaling downstream of NMBR (Figure 7A). This novel crosstalk is unique, to the best of our knowledge, as compared with a large body of studies on discrete GPCR crosstalk, which give rise to unique signaling output distinct from their respective canonical signaling pathway (71, 72). In the context of itch transmission, we have previously shown that mouse GRPR is able to cross talk with a number of GPCRs of unrelated families, including MOR1D, or human MOR1Y, 5HT1A and KOR, to regulate or modulate discrete types of itch transmission (49-51, 73). To the best of our knowledge, NPRC-NMBR cross-signaling represents the first example of the crosstalk between a GPCR and a non-GPCR receptor, the transmembrane guanylyl cyclase receptor. Although the mechanisms enabling G_αi/βγ_ signaling to amplify NMBR-mediated PLC-Ca^2+^ signaling are currently unknown, one can envision that such non-canonical cross-talk may confer spinal itch-specific neurons with much greater capability in coding complex behaviors.

### Respective Roles of BNP, SST and BNP/SST Fibers in Itch and Pain

While most human chronic itch conditions are resistant to antihistamines, existing mouse models of chronic itch, including allergic contact dermatitis (ACD) and atopic dermatitis (AD), are in part mediated by histamine-dependent mechanisms, in addition to histamine-independent mechanisms, because they are chemically, or haptens, induced allergic responses that involve histamine release from activated mast cells to mediate skin inflammation (6, 74). By contrast, dry skin itch in mice is mediated by GRP-dependent nonhistaminergic mechanism (21, 60, 75). Consistently, *Nppb* is upregulated in DRGs of mice with chronic itch comprising histaminergic itch signaling (22), whereas downregulated in dry skin itch, which is normal in *Npr1* KO mice.

In contrast to cutaneous GRP fibers (17), opto-stimulation of cutaneous *Nppb/Sst* fibers failed to induce scratching behavior, indicating that these fibers are not itch-specific. Indeed, our studies indicate that evoked scratching behavior by i.t. SST reflects both itch and pain components, suggesting concurrent disinhibition of itch and pain transmission by inhibition of SST2R GABAergic neural circuits. However, i.t. induced itch-related scratching behavior could be a pharmacological event that does not reflect the endogenous function of SST, similar to i.t. kappa opioid receptor agonists-evoked itch inhibition (50). There is some evidence supporting this possibility. First, conditional deletion of *Sst* or ablation of *Sst* neurons in DRGs does not impair itch transmission (31, 61). By contrast, these CKO mice exhibited enhanced thermal and mechanical pain (31). These data indicate that SST in sensory neurons is an inhibitory transmitter for pain rather than for itch (Figure S5). Although i.t. SST may evoke some scratching behavior related to itch, it is likely to reflect a pharmacological event, akin to morphine-induced pruritus. Consistent with this interpretation, a dramatic downregulation of *Sst* in DRGs of mice with chronic itch could be consequent to the overall dampening of spinal GABAergic neurons, as manifested in dramatically reduced expression of prodynorphin (50), which is expressed in some of the dorsal horn inhibitory neurons. Thus, optogenetic activation of cutaneous *Sst/Nppb/Nmb* fibers could result in inhibition of nociceptive transmission, which at the behavioral level is difficult to assess with the optical stimulation approach. Huang et al observed scratching behavior evoked by constant 20 Hz 470 nm stimulation of trigeminal ganglia cell bodies expressing channel rhodopsin (ChR2) in Sst-Cre/Ai32 (*Sst*^ChR2^) mice (31), but the nature of these scratching behaviors remains unclear. By contrast, direct opto-stimulation on the skin is a convenient way to mimic the activation of the itch terminals by pruritogens, as shown by optical stimulation of GRP fibers (17). Given an apparent role of endogenous STT from primary afferents in pain disinhibition and dispensable function in itch, we conclude that BNP/STT fibers are unlikely to function as itch-specific pruriceptors.

In summary, the present study demonstrates a novel BNP-NPRC-NMBR cross-signaling in the modulation of itch transmission (Figure 7A). By revealing the unexpected role for NPRC rather than NPRA in facilitating NMB mediated histaminergic itch transmission (Supplemental Figure 5), we clarify the inconsistencies in the literature regarding the widely reported BNP-NPRA/GRP-GRPR signaling pathway in itch transmission and bring conflicting results in a coherent framework. Together with previous studies, we have delineated distinct modes of action for GRP, NMB and BNP in the coding of histaminergic and nonhistaminergic itch information (Figure 7B). Whether there exist neuropeptides that similarly facilitate GRP-mediated nonhistaminergic itch transmission remains to be seen. These results should also help to gain a better understanding of distinct types of neuropeptides that are differentially involved in pathological and long-lasting under various pathological conditions.

## Methods

### Animals

Male mice between 7 and 12 weeks of age were used for experiments. C57BL/6J mice were purchased from The Jackson Laboratory (http://jaxmice.jax.org/strain/013636.html). *Npr1* KO mice (44), *Grpr* KO mice (76), *Nmbr* KO mice (77), BRAF^Nav1.8^ mice (21), and their wild-type (WT) littermates were used. We cross *Sst*^Cre^ mice (78) with a flox-stop channel rhodopsin-eYFP (ChR2-eYFP) line (Ai32) (79) to generate mice with ChR2-eYFP expression in Sst neurons (*Sst*^ChR2^). All experiments were performed in accordance with the guidelines of the National Institutes of Health and the International Association for the Study of Pain and were approved by the Animal Studies Committee at Washington University School of Medicine.

### Drugs and reagents

The dose of drugs and injection routes are indicated in figure legends. ANP, BNP, CNP, SST, and OCT were supplied from GenScript USA Inc. (Piscataway, NJ). GRP_18-27_ and NMB were from Bachem (King of Prussia, PA). Histamine, chloroquine (CQ), *Npr1* siRNA, *Npr2* siRNA, *Npr3* siRNA, and universal negative control siRNA were purchased from Sigma (St. Louis, MO). BNP-saporin (BNP-sap) and blank-saporin were made by Advanced Targeting Systems. Pertussis toxin (PTX) and gallein were from R&D Systems.

### Behavioral tests

Behavioral tests were videotaped (HDR-CX190 camera, Sony) from a side angle. The videos were played back on the computer and the quantification of mice behaviors was done by persons who were blinded to the treatments and genotypes.

### Acute scratching behavior

Itch behaviors were performed as previously described (13, 14). Briefly, mice were given 30 min to acclimate to the plastic arenas (10 × 10.5 × 15 cm). Mice were then briefly removed from the chamber for drug injections. Injection volume was 10 μL for i.t. injection and 50 μL for i.d. injection. Doses of drugs are indicated in figure legends. The number of scratching response was counted for 30 min at 5 min intervals. One scratching bout is defined as a lifting of the hind limb towards the body and then a replacement of the limb back to the floor or the mouth, regardless of how many scratching strokes take place between those two movements. Scratching towards injection site was counted after i.d. injection and all scratching bouts were counted after i.t. injection.

### Chronic itch models

*Dry Skin (Xerosis)*: The dry skin model was set up as described (60, 75). Briefly, the nape of mice was shaved and a mixture of acetone and diethyl ether (1:1) was painted on the neck skin for 15 s, followed by 30 s of distilled water application (AEW). This regimen was administrated twice daily for 9 days. Spontaneous scratching behaviors were recorded for 60 min in the morning before AEW treatment. BRAF^Nav1.8^ mice were used as described previously (21)

### RNAscope ISH

RNAscope ISH was performed as described (43, 50). Briefly, mice were anesthetized with a ketamine/xylazine cocktail (ketamine, 100 mg/kg and xylazine, 15 mg/kg) and perfused intracardially with 0.01 M PBS, pH 7.4, and 4 % paraformaldehyde (PFA). The spinal cord was dissected, post-fixed in 4 % PFA for 16 h, and cryoprotected in 20% sucrose overnight at 4 °C. Tissues were subsequently cut into 18 μm-thick sections, adhered to Superfrost Plus slides (Fisher Scientific), and frozen at −20 °C. Samples were processed according to the manufacturer’s instructions in the RNAscope Fluorescent Multiplex Assay v2 manual for fixed frozen tissue (Advanced Cell Diagnostics), and coverslipped with Fluoromount-G antifade reagent (Southern Biotech) with DAPI (Molecular Probes). The following probes, purchased from Advanced Cell Diagnostics were used: *Nppb* (nucleotide target region 4 - 777; accession number NM_008726.5), *Sst* (nucleotide target region 18 - 407; accession number NM_009215.1), *Npr1* (nucleotide target region 941 - 1882; accession number NM_008727.5), *Npr2* (nucleotide target region 1162 - 2281; accession number NM_173788.3), *Npr3* (nucleotide target region 919 - 1888; accession number NM_008728.2), *Grp* (nucleotide target region 22 - 825; accession number NM_175012.2), *Grpr* (nucleotide target region 463-1596; accession number - NM_008177.2), and *Nmbr* (nucleotide target region 25 - 1131; accession number NM_008703.2). Sections were subsequently imaged on a Nikon C2+ confocal microscope (Nikon Instruments, Inc.) in three channels with a 20X objective lens. The criterion for including cells as positive for a gene expression detected by RNAscope ISH: A cell was included as positive if two punctate dots were present in the nucleus and/or cytoplasm. For co-localization studies, dots associated with single DAPI stained nuclei were assessed as being co-localized. Cell counting was done by a person who was blinded to the experimental design.

### ISH and immunohistochemistry

ISH was performed using digoxigenin-labeled cRNA probes as previously described (80). Briefly, mice were anesthetized with an overdose of a ketamine/xylazine cocktail and fixed by intracardiac perfusion of cold 0.01 M PBS, pH 7.4, and 4% paraformaldehyde. The spinal cord, DRG, and hairy nape skin tissues were immediately removed, post-fixed in the same fixative overnight at 4°C, and cryoprotected in 20% sucrose solution. DRGs, lumbar spinal regions, and hairy nape skin were frozen in OCT and sectioned at 20 μm thickness on a cryostat. Immunohistochemical (IHC) staining was performed as described (81). Spinal cord and DRG tissues were sectioned at 20 μm thickness. Hairy nape skin was sectioned at 30 μm thickness. Free floating sections were incubated in a blocking solution containing 2% donkey serum and 0.1% Triton X-100 in PBS (PBS-T) for 2 h at room temperature. The sections were incubated with primary antibodies overnight at 4°C, washed three times in PBS, incubated with the secondary antibodies for 2 h at room temperature and washed three times. Sections were mounted on slides with Fluoromount G (Southern Biotech) and coverslips. Fluorescein isothiocyanate (FITC)-conjugated Isolectin B4 from *Griffonia simplicifolia* (IB4, 10 μg/mL; L2895, Sigma), IB4-AlexaFluor 568 conjugate (2 μg/mL, ThermoFisher Scientific) or the following primary antibodies were used: rabbit anti-CGRPα (1:3000; AB1971, Millipore), guinea pig anti-Substance P (1:1000; ab10353, Abcam), guinea pig anti-TRPV1 (1:1000; GP14100, Neuromics), chicken anti-NF-H (1:2000, EnCor Biotechnology, CPCA-NF-H), rabbit anti-βIII-Tubulin (1:2000, Biolegend, 802001), rabbit anti-GFP (1:1000, Molecular Probes, A11122), chicken anti-GFP (1:500, Aves Labs, GFP-1020). The secondary antibodies were purchased from Jackson ImmunoResearch Laboratories including Cyanine 3 (Cy3), Cyanine 5 (Cy5) - or FITC conjugated donkey anti-rabbit, anti-mouse, anti-chicken or anti-guinea pig IgG (Cy3, 0.5 μg/ml; FITC, 1.25 μg/mL), biotin-SP (long-spacer)-conjugated donkey anti-rabbit IgG (1 μg/mL) and avidin-conjugated Alexa Fluor 488 (0.33 μg/mL). Images were taken using a Nikon Eclipse Ti-U microscope with Cool Snap HQ Fluorescent Camera and DS-U3 Brightfield Camera controlled by Nikon Elements Software (Nikon) or a Leica TCS SPE confocal microscope with Leica LAS AF Software (Leica Microsystems). The staining was quantified by a person blinded to the genotype using ImageJ (version 1.34e, NIH Image) as previously described (21). Images were taken using a Nikon C2+ confocal microscope system (Nikon Instruments, Inc.) and analysis of images was performed using ImageJ software from NIH Image (version 1.34e). At least three mice per group and 10 lumbar sections across each group were included for statistical comparison.

### Small interfering RNA treatment

Negative control siRNA (SIC001) and selective siRNA duplex for mouse *Npr1* (SASI_Mm01_00106966), mouse *Npr2* (SASI_Mm01_00201357), and mouse *Npr3* (SASI_Mm01_00036567) were purchased from Sigma. RNA was dissolved in diethyl pyrocarbonate-treated PBS and prepared immediately prior to administration by mixing the RNA solution with a transfection reagent, RVG-9R (Genscript). The final concentration of RNA was 2 μg/10 μL. siRNA was delivered to the lumbar region of the spinal cord. The injections were given once daily for 6 consecutive days as described previously with some modifications (51, 82, 83). Behavior testing was carried out 24 h after the last injection.

### BNP-saporin treatment

Mice were treated with one-time i.t. injection of BNP-sap (2.5 μg/mouse) as previously described (25) with the reduced dose, due to the lethal effect of BNP-sap (5 μg). Behavioral tests were performed 2 weeks after BNP-sap injection. After behavioral tests, the spinal cord/DRGs of mice were processed for real-time RT-PCR and ISH.

### Real-time RT-PCR

Real-time RT-PCR was performed as previously described with Fast-Start Universal SYBR Green Master (Roche Applied Science) (35, 51). All samples were assayed in duplicate (heating at 95°C for 10 s and at 60°C for 30 s). Data were analyzed using the Comparative CT Method (StepOne Software version 2.2.2.), and the expression of target mRNA was normalized to the expression of *Actb* and *Gapdh*. The following primers were used: *Nppb* (NM_008726.4): 5’- GTCAGTCGTTTGGGCTGTAAC-3’, 5’- AGACCCAGGCAGAGTCAGAA-3’; amplicon size, 89 bp; Sst (NM_009215.1): 5’− CCCAGACTCCGTCAGTTTCT −3’, 5’− CAGCAGCTCTGCCAAGAAGT −3’, amplicon size, 87 bp; *Npr1* (NM_008727.5): 5’- TGGAGACACAGTCAACACAGC-3’, 5’- CGAAGACAAGTGGATCCTGAG-3’; amplicon size, 70 bp; *Npr2* (NM_173788.3): 5’- TGAGCAAGCCACCCACTT-3’, 5’- AGGGGGCCGCAGATATAC-3’, amplicon size, 60 bp; *Npr3* (NM_008728.2): 5’- TGCACACGTCTGCCTACAAT-3’, 5’- GCACCGCCAACATGATTCTC -3’, amplicon size, 138 bp; *Grpr* (NM_008177.2): 5’- TGATTCAGAGTGCCTACAATCTTC-3’, 5’- TTCCGGGATTCGATCTG -3’; amplicon size, 71 bp; *Nmbr* (NM_008703.2): 5’- GGGGGTTTCTGTGTTCACTC -3’, 5’- CATGGGGTTCACGATAGCTC -3’, amplicon size, 67 bp; *Actb* (NM_007393.3): 5’- TGTTACCAACTGGGACGACA-3’, 5’- GGGGTGTTGAAGGTCTCAAA-3’; amplicon size, 166 bp; and *Gapdh* (NM_008084.2): 5’-CCCAGCAAGGACACTGAGCAA-3’, 5’- TTATGGGGGTCTGGGATGGAAA-3’; amplicon size, 93 bp.

### Dissociation of spinal neurons and calcium imaging

Primary culture of spinal dorsal horn neurons was prepared from 5 ~ 7 day old C57BL/6J mice (Zhao et al., 2014b). After decapitation, laminectomy was performed and the dorsal horn of the spinal cord was dissected out with a razor blade and incubated in Neurobasal-A Medium (Gibco) containing 30 μL papain (Worthington) at 37°C for 20 min. Enzymatic digestion was stopped by replacing it with a 1 ml Neurobasal-A medium. After washing with the same medium three times, gentle trituration was performed using a flame polished glass pipette until the solution became cloudy. The homogenate was centrifuged at 1,500 rpm for 5 min and the supernatant was discarded. The cell pellet was re-suspended in culture medium composed of Neurobasal-A medium (Gibco, 92% vol/vol), fetal bovine serum (Invitrogen, 2% vol/vol), HI Horse Serum (Invitrogen, 2% vol/vol), GlutaMax (2 mM, Invitrogen, 1% vol/vol), B27 (Invitrogen, 2% vol/vol), Penicillin (100 μg/mL) and Streptomycin (100 μg/mL) and plated onto 12-mm coverslips coated with poly-D-lysine. After three days in culture, neurons were used for calcium imaging as described previously (Zhao et al., 2014b). Neurons were loaded with Fura 2-acetomethoxy ester (Molecular Probes) for 30 min at 37°C. After washing, neurons were imaged at 340 and 380 nm excitation to detect intracellular free calcium.

### Optical skin stimulation behavior

*Sst*^ChR2^ mice and wild-type littermates (*Sst*^WT^) were used for optical skin stimulation experiments. The nape skin was shaved 3 days prior to stimulation in all mice tested. One day prior to the experiments, each mouse was placed in a plastic arena (10 × 11 × 15 cm) for 30 min to acclimate. For blue light skin stimulation, a fiber optic cable was attached to a fiber-coupled 473 nm blue laser (BL473T8-150FC, Shanghai Laser and Optics Co.) with an ADR-800A adjustable power supply. Laser power output from the fiber optic cable was measured using a photometer (Thor Labs) and set to 15 mW from the fiber tip. An Arduino UNO Rev 3 circuit board (Arduino) was programmed and attached to the laser via a BNC input to control the frequency and timing of the stimulation (1, 5, 10 or 20Hz with 10 ms on-pulse and 3 s On – 3 s Off cycle for 5 min). During stimulation, the mouse was traced manually by a fiber optic cable with a ferrule tip that was placed 1-2 cm above the nape skin. Videos were played back on a computer for scratching behavior assessments by observers blinded to the animal groups and genotypes.

### Statistical analyses

Values are reported as the mean ± standard error of the mean (SEM). Statistical analyses were performed using Prism 6 (v6.0e, GraphPad, San Diego, CA). For comparison between two or more groups, unpaired, paired two-tailed t-test, one-way ANOVA followed by Tukey post hoc tests, or two-way repeated measures ANOVA followed by Sidak’s post hoc analysis, was used. Normality and equal variance tests were performed for all statistical analyses. p < 0.05 was considered statistically significant.

## Author contributions

Q.T.M., X.Y.L., X.T.L., D.M.B. and Z.F.C. designed the experiments and Q.T.M. and H.J performed behavioral experiments. X.Y.L. performed molecular analysis, X. T. L., D.M.B. and Q. Y. Y. performed in situ hybridization and immunohistochemistry. S. Y., L.W., A. M., J. H. J., R. K., K.F. S., and J.Y. contributed to the project. X.Y.L. and Z.F.C wrote the manuscript and Q.T.M and D.M.B. participated in the writing. Q.T.M., X.Y.L., and X. T. L., contributed equally to this work.

## Acknowledgements

We thank N. Maeda for *Npr1*^+/−^ mice, TRC, CDI, and the TGI for RNAi plasmids, and the Chen lab members for comments. The project has been supported by the National Science and Technology Major Project of China (2016ZX08011-005) (A. T) and the NIH grants 1R01AR056318-06, R21 NS088861-01A1, R01NS094344, R01 DA037261-01A1 and R56 AR064294-01A1 (Z. F. C).

## SUPPLEMENTAL FIGURES AND FIGURE LEGENDS

**Figure S1.**
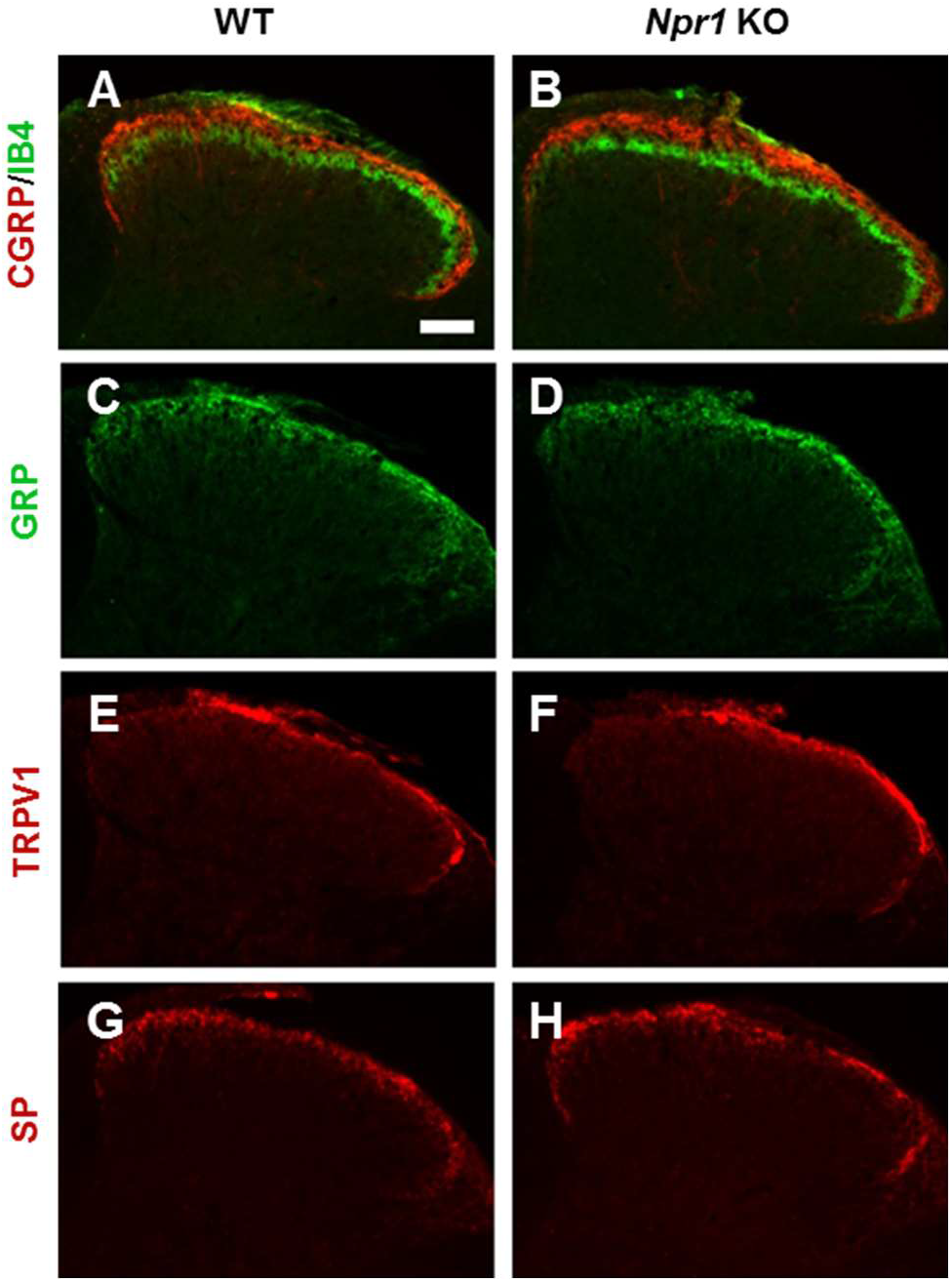
Normal innervation of primary afferents in *Npr1* KO mice and WT mice. (**A** and **B**) Density and innervation of CGRP^+^ (red) and IB4^+^ fibers (green) in the superficial dorsal horn are comparable in WT (**A**) and *Npr1* KO mice (**B**). (**C-H**) GRP^+^, TRPV1^+^, and SP^+^ primary afferents in the superficial dorsal horn.

**Figure S2.**
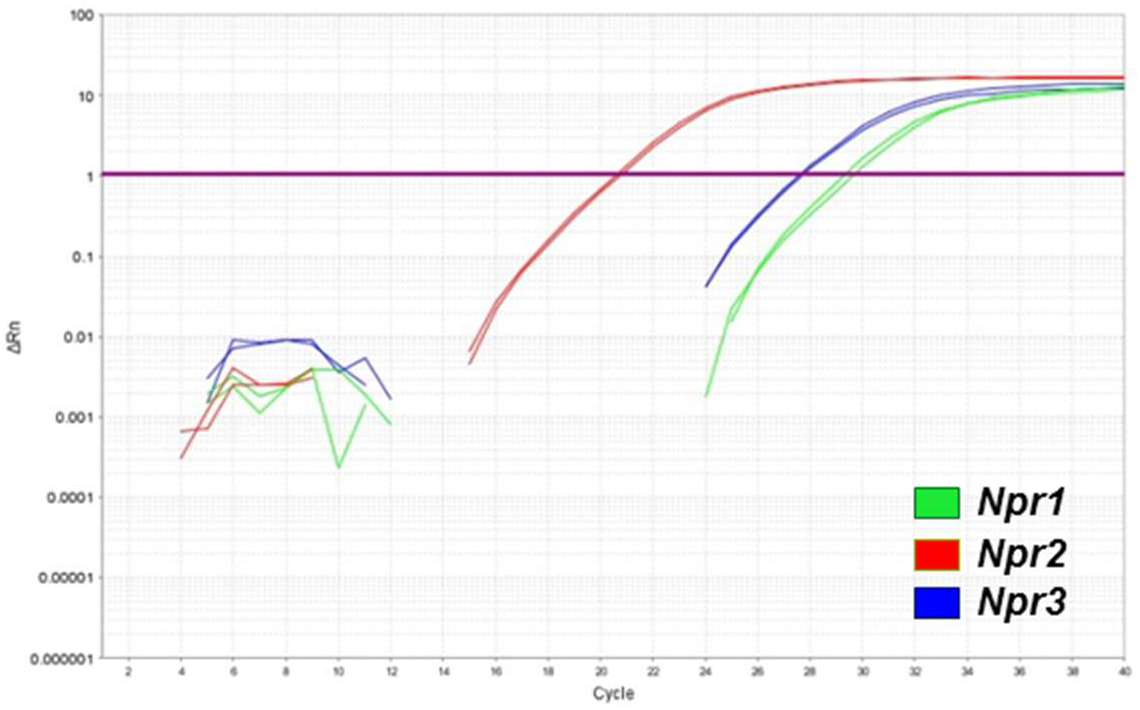
*Npr1*, *Npr2*, and *Npr3* are endogenously expressed in HEK 293 cells as detected by real-time RT-PCR.

**Figure S3.**
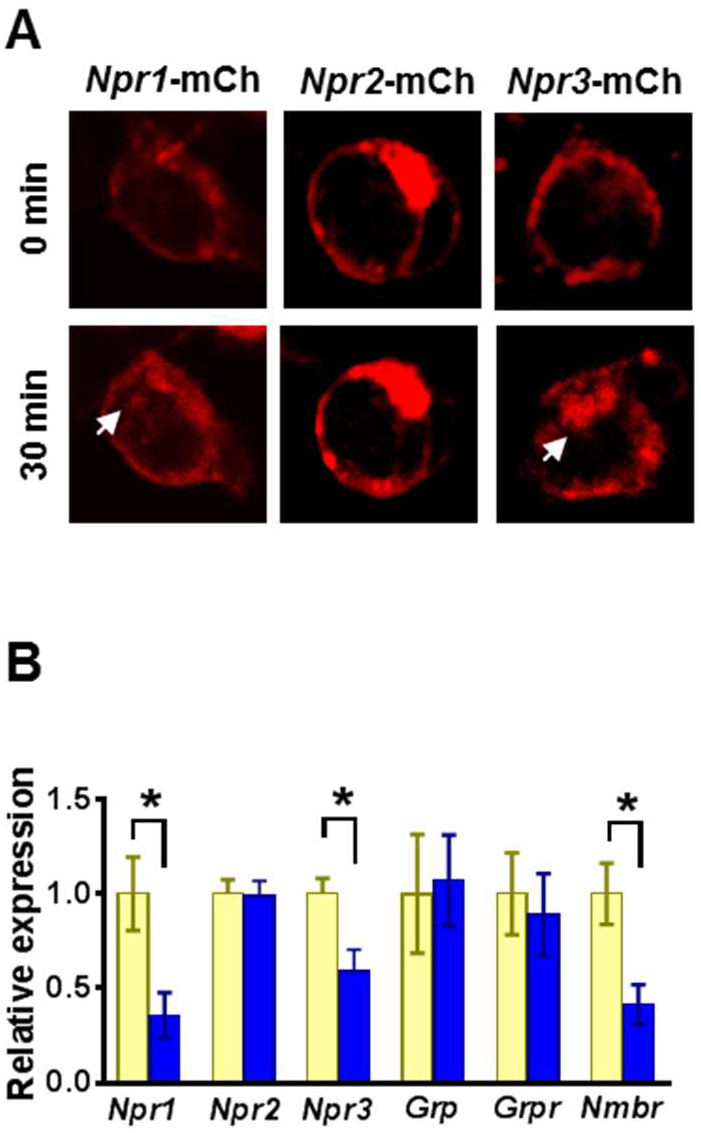
BNP induces internalization of *Npr1* and *Npr3*. (**A**) Incubation of BNP (10 μM) for 30 min caused internalization of *Npr1*-mCh and *Npr3*-mCh as indicated by arrows. Scale bar, 20 μm. (**B**) Real-time RT-PCR results show that *Npr1*, *Npr3* and *Nmbr* mRNA levels were significantly reduced in the dorsal spinal cord of BNP-sap treated mice. *Npr2* and *Grpr* mRNA levels were not affected by BNP-sap treatment. n = 4. \**P* < 0.05, unpaired t test.

**Figure S4.**
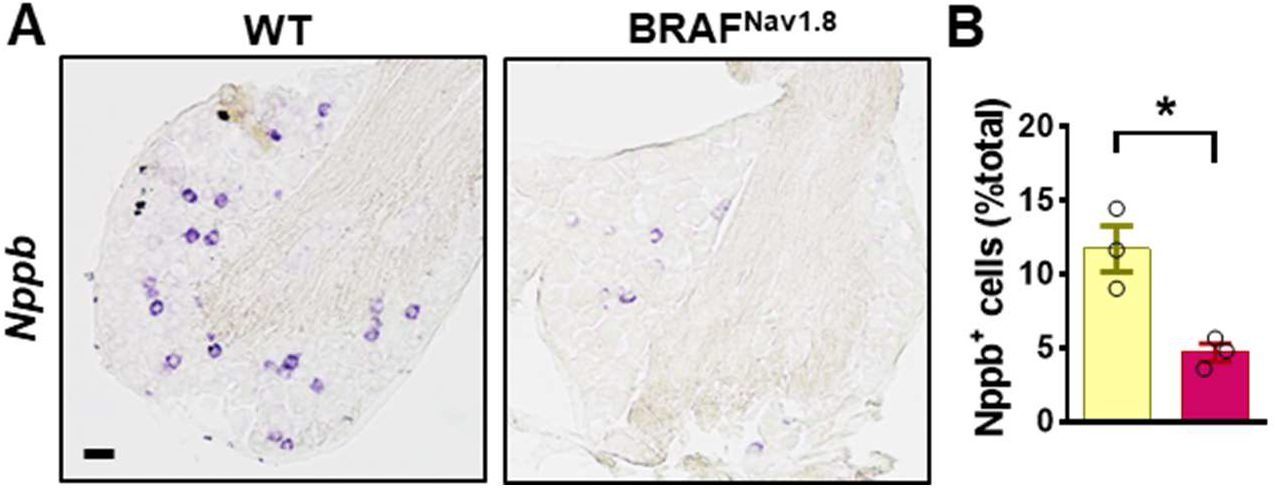
BNP-NPRA signaling is dispensable for chronic itch. (**A** and **B**) ISH of *Nppb*, in DRGs of WT mice and BRAF^Nav1.8^ mice. n = 4. Values are presented as mean ± SEM. Scale bars, 50 μm

**Figure S5.**
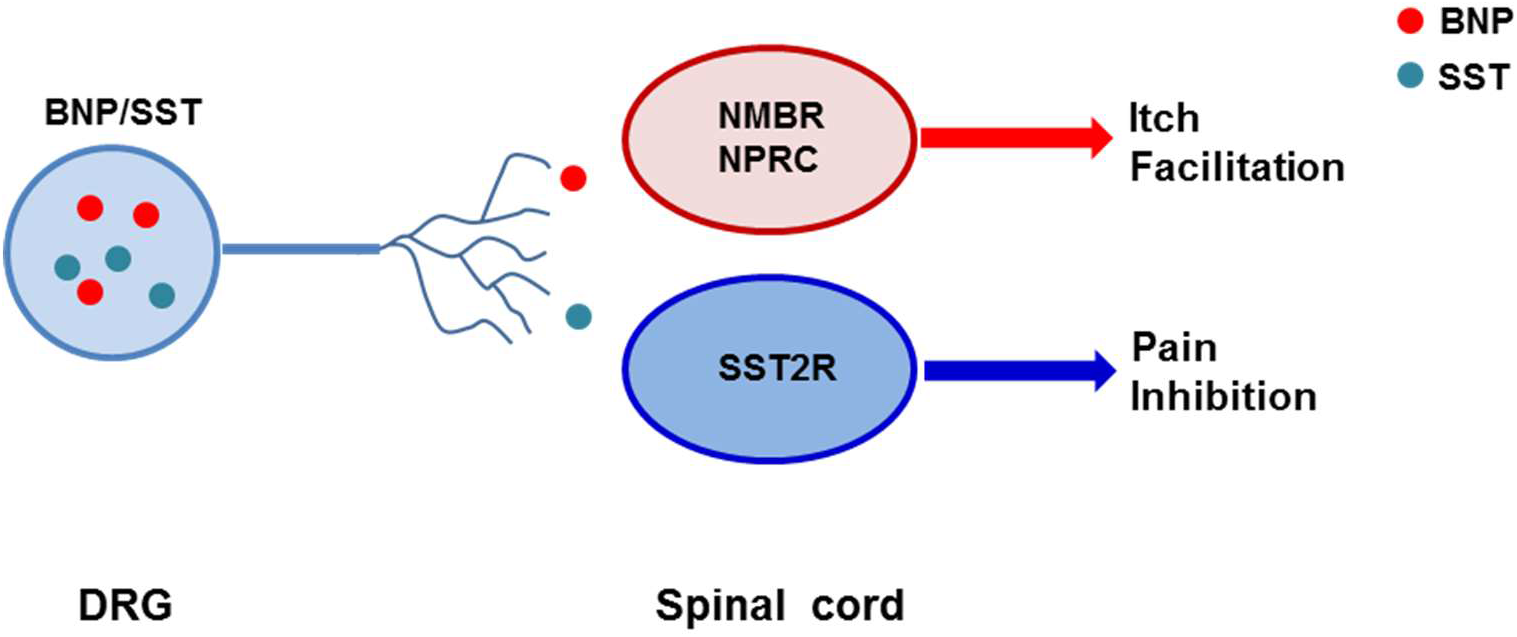
A hypothetic model depicting the role of BNP and SST in itch facilitation and pain inhibition, respectively.

